# The role of environment, local adaptation and past climate fluctuation on the amount and distribution of genetic diversity in the teosinte in Mexico

**DOI:** 10.1101/820126

**Authors:** Jaime Gasca-Pineda, Yocelyn T. Gutiérrez-Guerrero, Erika Aguirre-Planter, Luis E. Eguiarte

## Abstract

Wild maize, commonly known as teosinte, has a wide distribution in central Mexico and inhabits a wide range of environmental conditions. According to previous studies, the environment is a determinant factor for the amount and distribution of genetic diversity. In this study, we used a set of neutral markers to explore the influence of contemporary factors and historical environmental shifts on genetic diversity, including present and three historical periods. Using a set of 22 nuclear microsatellite loci, we genotyped 527 individuals from 29 localities. We found highly variable levels of genetic diversity (*Z. m. parviglumis H*_*E*_= 0.3646–0.7699; *Z. m. mexicana H*_*E*_= 0.5885–0.7671) and significant genetic structure among localities (average *D*_*EST*_= 0.4332). Also, we recovered significant values of heterozygote deficiency (average *F*_*IS*_= 0.1796) and variable levels of selfing (*sg*^*2*^=0.0–0.3090). The Bayesian assignment analysis yielded four genetic clusters dividing the sample into subspecies, that in turn, were separated into two clusters. Environmental conditions played a strong influence in the distribution of genetic diversity, as demographic analysis and changes in species range revealed by modeling analyses were consistent. We conclude that current genetic diversity in teosinte is the result of a mixture of local adaptation and genetic isolation along with historical environmental fluctuations.

## Introduction

*Zea mays* comprises the cultivated maize (*Zea mays* ssp. mays L.), as well as wild maize populations commonly known as teosintes (Doebley & Iltis, 1980). The most common teosinte subspecies, *Z. m. parviglumis* and *Z. m. mexicana*, diverged ca. 61,000 years ago, and both are abundant and widely distributed in central-west México (Aguirre-Liguori, Aguirre-Planter & Eguiarte, 2016; Hufford *et al.*, 2012). *Z. m. parviglumis* grows in the lowlands (<1900m above sea level) in warm and moist conditions, while *Z. m. mexicana* is found in colder and dryer highlands (>1500m above sea level) (Aguirre-Liguori, Aguirre-Planter & Eguiarte, 2016). Aside from the intrinsic interest due to their relationship with cultivated crops (Aguirre-Liguori, Aguirre-Planter & Eguiarte, 2016), the wide distribution of teosintes and the contrasting environmental conditions inhabited by each subspecies make them an attractive system to study microevolutionary processes, including genetic divergence, local adaptation, and phylogeographic patterns. Moreover, the public resources available for *Zea mays* (for example, https://www.maizegdb.org), allow selecting genetic markers to avoid genetic-physical linkage, and in addition, a number of loci have already been validated in studies of genetic diversity in maize (Fukunaga *et al.*, 2005; Orozco-Ramírez *et al.*, 2016).

Previous studies dealing with teosintes genetic diversity were focused on genetic resources and domestication processes (Fukunaga *et al.*, 2005; van Heerwaarden *et al.*, 2010), evolutionary analyses based on particular genes (Moeller, Tenaillon & Tiffin, 2007; Ross-Ibarra, Tenaillon & Gaut, 2009), and genomic adaptation (Fustier *et al.*, 2017; Aguirre-Liguori *et al.*, 2017; 2019). However, due to their specific objectives, the majority of these studies used broad geographic scales or a large number of localities with few individuals, while other studies involved a higher number of individuals, yet using few molecular markers, either isozyme or microsatellites (Buckler, Gaut & McMullen, 2006; Fukunga *et al.*, 2005). More recently, some studies implemented SNPs (Pyhäjärvi *et al.*, 2013; Aguirre-Liguori *et al.*, 2017). In general, these studies suggested that genetic diversity and the distribution of teosintes are strongly affected by environmental factors (Aguirre-Liguori *et al.*, 2017). Thus, it is expected that historical events like the Last Glacial Maximum (LGM) and the Younger dryas (Berger, 1990) influenced the distribution of teosinte populations, with a concurrent influence on their genetic diversity. Moreover, aside from the environmental factors shaping teosinte genetic diversity, previous studies left open questions about the fine details of microevolution, such as genetic structure, past demographic events, selfing rates, and the geographical distribution of genetic diversity. Hence, relevant questions have remained: Are there high levels of genetic structure among teosinte populations, or they comprise a genetic continuum? Is the amount and distribution of current genetic diversity influenced by the past climate changes? Are local factors, like selfing and local adaptation shaping genetic diversity? And, using neutral molecular data, is there evidence of gene flow between subspecies?

Recent advances in coalescent theory (Wu & Drummond, 2011; Nikolic & Chevalet, 2014) used jointly with distribution modeling analyses for present and past environmental conditions (Hijmans *et al.*, 2005; Thuiller *et al.*, 2019), have proved to be powerful tools to understand the processes underlying the observed levels and patterns of genetic variation in plants and animals (Aguirre-Liguori *et al.*, 2019; Castellanos-Morales *et al.*, 2019). In this sense, neutral molecular markers, such as microsatellites allow for a wide variety of robust analyses (Selkoe & Toonen, 2006; Guichoux *et al.*, 2011; Kalia *et al.*, 2011) including the inference of demographics and migration process. Even though the many appealing analyses that can be performed using microsatellites, recent studies highlighted the need of a high number of loci (>20) along with a comprehensive geographic representation of the species (including large number of populations and individuals), in order to achieve reliable conclusions (Aguirre-Liguori *et al.*, submitted).

In this study, we used a set of 22 microsatellite loci to conduct thorough analyses of the amount and geographic distribution of genetic diversity for a total of 29 teosinte populations in Mexico, 14 of *Z. m. parviglumis* and 15 of *Z. m. mexicana*. The studied localities encompassed a wide range of environmental conditions and most of the geographic distribution of both subspecies (Aguirre-Liguori *et al.*, 2017). We estimated the genetic diversity, genetic structure, inbreeding and selfing levels, and identified genetic barriers among 29 localities of teosinte. Additionally, we used present and past environmental data to assess the influence of past environmental conditions in the demographic history of this species.

## Materials and methods

### Sampling, DNA extraction and genotyping

We sampled 29 sites including 14 localities of *Zea mays parviglumis* and 15 localities of *Zea mays mexicana* subspecies (Fig. 1, Table S1, Supplementary Materials) encompassing a broad altitudinal and environmental gradient of teosinte populations. Seeds were collected from 10-26 plants per site, and one seed per plant was grown in a greenhouse obtaining a total of 527 plants. DNA extraction was carried out from fresh leaf tissue using a modified version of the CTAB extraction method (Doyle & Doyle, 1987).

**Figure 1.**
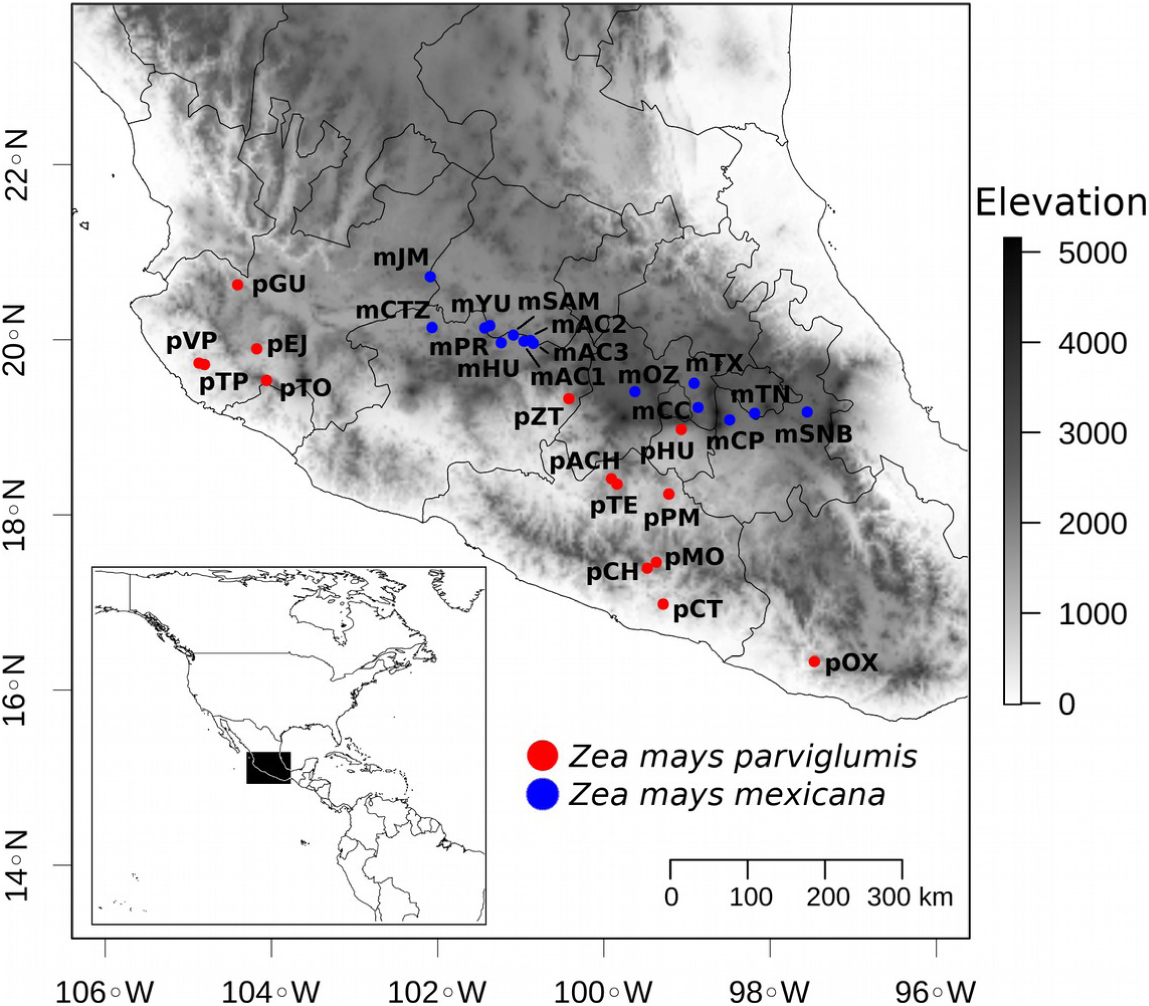
Map depicting teosinte sampled localities. In red *Zea mays parviglumis*, in blue *Zea mays mexicana*.

To evaluate the genetic diversity of teosintes, we amplified a set of 22 microsatellite loci previously used for maize genetic diversity analyses. To avoid physical linkage, loci were selected considering their chromosomal position and distance within chromosomes (>150cM). The microsatellites names and primer sequences are reported in Supplementary Materials (Table S2). The PCR products were visualized on 2% agarose gels stained with ethidium bromide. The fluorescent-labeled PCR products were genotyped in the UIUC Core Sequencing Facility (https://unicorn.biotech.illinois.edu/) using GeneScan LIZ 600 size standard (Applied Biosystems). Allele sizes were determined using Genemapper v 3.7 (Applied Biosystems). Presence of genotyping errors and null alleles were assessed using MICRO-CHECKER v 2.2.3 (Van Oosterhout *et al.*, 2004), applying the Bonferroni correction for multiple comparisons. To evaluate potential bias due to null alleles in estimates of genetic structure, we used the FreeNa software (Chapuis & Estoup, 2007) to implement a statistical test for differences between the standard *F*_*ST*_ versus the *F*_*ST*_ with the ENA (Excluding Null Alleles) correction.

### Population Genetics Analyses

We estimated basic population genetics summary statistics: observed number of alleles (*Na*), the rarefied allelic richness (*Ar*, rarefied to the minimum sample size), the observed (*H*_*O*_) and expected heterozygosity (*H*_*E*_), using the package “adegenet” version 2.1.1 (Jombart, 2008) implemented in R (R Core Team, 2015).

To detect deviations from Hardy-Weinberg proportions, we performed a test of excess or deficiency of heterozygotes by means the fixation index (*F*_*IS*_) (Wright, 1965; Weir & Cockerham, 1984) using Genepop version 4.6 (Rousset, 2008). Its significance was calculated using the Fisher exact test with 10,000 dememorisation steps, 1,000 batches, and 5,000 iterations per batch. To approximate the degree of selfing in teosinte localities, we calculated *sg*^*2*^ based on multilocus comparison of linkage disequilibria using the software RMES (David *et al.*, 2007).

To evaluate the genetic structure among localities, we used the R package DEMEtics (Gerlach *et al.*, 2010) to estimate the Jost’s *D*_*EST*_. We selected this measure because standard *F*_*ST*_ may underestimate population differentiation due to the high mutation rates of microsatellites (Balloux & Lugon-Moulin, 2002), and large population sizes of teosintes. We calculated the 95% confidence intervals (C.I.) as a measure of significance; if *D*_*EST*_ C.I. values did not approximate zero, we took the value as significant. Additionally, we performed a Principal Component Analysis (PCA) to evaluate the allelic differences at the individual level using the sampling localities as point classes with “adegenet” R package.

To detect genetic grouping among individuals and localities, we ran Structure v2.3 (Pritchard, Stephens & Donnelly, 2000) using the admixture model with correlated allele frequencies (Falush, Stephens & Pritchard, 2003). We performed 25 independent runs for each value of *K* from 1 to 30 using 1,000,000 MCMC steps followed by 500,000 steps as burn-in. To find the most likely number of clusters (*K*) we used the method proposed by Evanno, Regnaut & Goudet (2005) implemented in Structure Harvester (Earl & vonHoldt, 2012). Also, all lnP(D) values were plotted to choose the *K* value with the highest likelihood and lowest variance. We performed the analyses for the whole sample, as well for each subspecies alone.

Gene flow was approximated by means of the recent migration rates (*m*) estimated by BayesASS 3.0.3 (Wilson & Rannala, 2003). As BayesASS does not perform well with multiple gene pools within populations (Wilson & Rannala, 2003), we used the genetic clusters obtained by Structure. To ensure good chain-mixing, we performed several test runs varying the values of parameters to achieve acceptance rations of 0.25-60 as recommended by BayesASS authors. The analysis was performed using 10×10^9^ Markov chain Monte Carlo iterations, sampling every 100,000 steps and discarding the first 10% iterations as burn-in period.

In order to detect genetic barriers among localities, we implemented the Monmonier algorithm (Monmonier, 1973) using adegenet. To perform this analysis, we used a Gabriel graph as a connection network among localities and used the localities’ scores obtained from the previously estimated PCA as distance matrix. The distance threshold between immediate neighbors was chosen as an abrupt decrease between connected points. We used the function optimize.monmonier to obtain the best boundaries.

### Isolation by Distance and Isolation by Environment analyses

To approximate Isolation by Distance (IBD) patterns for each subspecies, we performed standard Mantel tests (Mantel, 1967) using pairwise Nei genetic distances between localities versus the log10 of geographic distances in meters. The Mantel tests and their significance (10,000 random replications) were estimated by means of the R package vegan (Oksanen *et al.*, 2009), while geographic distances were obtained using the raster package (Hijmans, 2019).

We evaluated the relative effects of geographical distance and environment on genetic differentiation using R package Sunder (Botta *et al.*, 2014). This method uses the decay in covariance in allele frequencies between populations as a function of the geographical and ecological distances (here, the environmental distance matrix). We implemented three models, *G*, where geographic distance alone explains the decline in covariance in allele frequencies; *E*, where environmental distance explains the decline; and *G+E*, where both distances explain equally the decline in the allele frequencies. We ran the analysis using 10 million iterations, sampling every 1,000 iterations. The initial state and upper bounds of the Dirichlet prior parameters distribution were implemented as suggested by the software authors. The convergence was assessed by plots of parameters’ traces, and we used 10 % of the data as a validation set for the cross-validation step. The environmental distance matrix was generated using euclidean distances estimated from the bioclim dataset obtained for the SDM analyses (see below).

### Demographic analyses

We approximated changes in the effective population size (*N*_*e*_) of teosinte populations using R package VarEff v1.2 (Nikolic & Chevalet, 2014). This package estimates the effective population sizes from present to ancestral time by the simulation of demographic steps of constant size using a coalescent approach. The model calculates posterior probabilities for each step through likelihood approximation and plots the results in terms of *N*_*e*_ through time. We used previously reported microsatellite mutation rate range for teosinte (average=7.7×10^−4^ per generation, 95% HPD 1.1×10^−3^−5.2×10^−4^) (Vigouroux *et al.*, 2002) to approximate the time in generations for the fluctuation on *N*_*e*_. For demographic analyses, we used the genetic clusters obtained with Structure. We ran the program for 100,000 steps with a batch length of 10 steps and sampling every 10 steps with an acceptance ratio of 0.25. We used the Two-Phase model (Di Rienzo *et al.*, 1994) with the proportion of mutational events (*c*) fixed at 0.15. As both subspecies are recognized as annual plants (Aguirre-Liguori, Aguirre-Planter & Eguiarte, 2016) we considered one year as generation time.

### Spatial Distribution Modeling

We gathered a total of 375 georeferenced points including this study and previous published records of *Z. m. parviglumis* and *Z. m. mexicana* (www.conabio.org). In order to reduce bias due to autocorrelation caused by local overrepresentation, we generated a 1/10 degree grid and then selected one random point using the R package “raster”. The final data set included 62 points for *Z. m. mexicana* and 69 points for *Z. m. parviglumis*. We obtained 19 current environmental layers from the Worldclim database (http://worldclim.org/bioclim, Hijmans *et al.*, 2005). To evaluate the influence of past climate conditions in teosinte distribution, we included the past Bioclimatic variables for the Mid-Holocene (MID), Last Glacial Maximum (LGM), and Last Interglacial (LIG) (Otto-Bliesner *et al.*, 2006). To reduce variable redundancy, we conducted a paired Spearman’s rank correlation test and excluded variables with values >|0.75|. The final set included the following ten variables: BIO2 (Mean Diurnal Range), BIO3 (Isothermality), BIO4 (Temperature Seasonality), BIO8 (Mean Temperature of Wettest Quarter), BIO12 (Annual Precipitation), BIO14 (Precipitation of Driest Month), BIO15 (Precipitation Seasonality), BIO17 (Precipitation of Driest Quarter), BIO18 (Precipitation of Warmest Quarter), BIO19 (Precipitation of Coldest Quarter).

To generate the Species Distribution Models (SDMs) we used the platform biomod2 v 3.3 (Thuiller *et al.*, 2019) in R. The SDMs were constructed by means of ensemble forecasting models (Araújo & New, 2007), using the committee-averaging criteria. We implemented four modelling algorithms in our analysis: 1) Generalised Linear Model (GLM; McCullagh & Nelder, 1989); 2) Generalized Boosted Model (GBM; Friedman 1991); 3) MaxEnt (Phillips, Anderson & Schapire, 2006); and 5) Random Forest (RF; Breiman, 2001). Occurrence data was split into 70% for model calibration and 30% for model evaluation. Using this 70%-30% criteria, five random-sampled data sets were run for each algorithm. Then, we generated five pseudo-absence datasets of 5,000 points selected at random. Model performance was assessed using the true skill statistic (TSS) and the area under the receiver operating characteristic curve (AUC), and models that did not meet thresholds of TSS>0.8 and 0.95>AUC were discarded. Final ensembles were transformed to binary points selecting the threshold that maximized the TSS score.

## Results

Overall, we detected a total of 355 alleles for the 22 microsatellite loci analyzed with an allelic range from six to 30 alleles per locus (allelic frequencies for all individuals and markers are reported in Table S3). The overall genetic diversity was *H*_*E*_= 0.818 (s.d.= 0.077) and *H*_*O*_= 0.565 (s.d.= 0.104) (values reported in Table S4). At subspecies level, *Z. m. parviglumis* showed an average *H*_*E*_= 0.666 (s.d. = 0.112), *H*_*O*_= 0.538 (s.d. = 0.103), and allelic richness, *Ar*= 3.948 (s.d.= 0.683), while *Z. m. mexicana* had *H*_*E*_= 0.714 (s.d. = 0.047), *H*_*O*_= 0.585 (s.d=0.044), and *Ar*= 4.198 (s.d.=0.760). Furthermore, we found a wide range of genetic diversity among localities, ranging from *H*_*E*_= 0.364 (s.d. = 0.172) to *H*_*E*_= 0.769 (s.d. = 0.115), and an allelic richness from *Ar*= 2.335 (s.d. = 1.172) to *Ar*= 4.749 (s.d.= 1.218) for pVP and pCH respectively (Fig. 2a, Table S4). We did not find a statistical correlation between genetic diversity and sample size per locality (*r*^*2*^<0.01, *p*>0.2).

**Figure 2.**
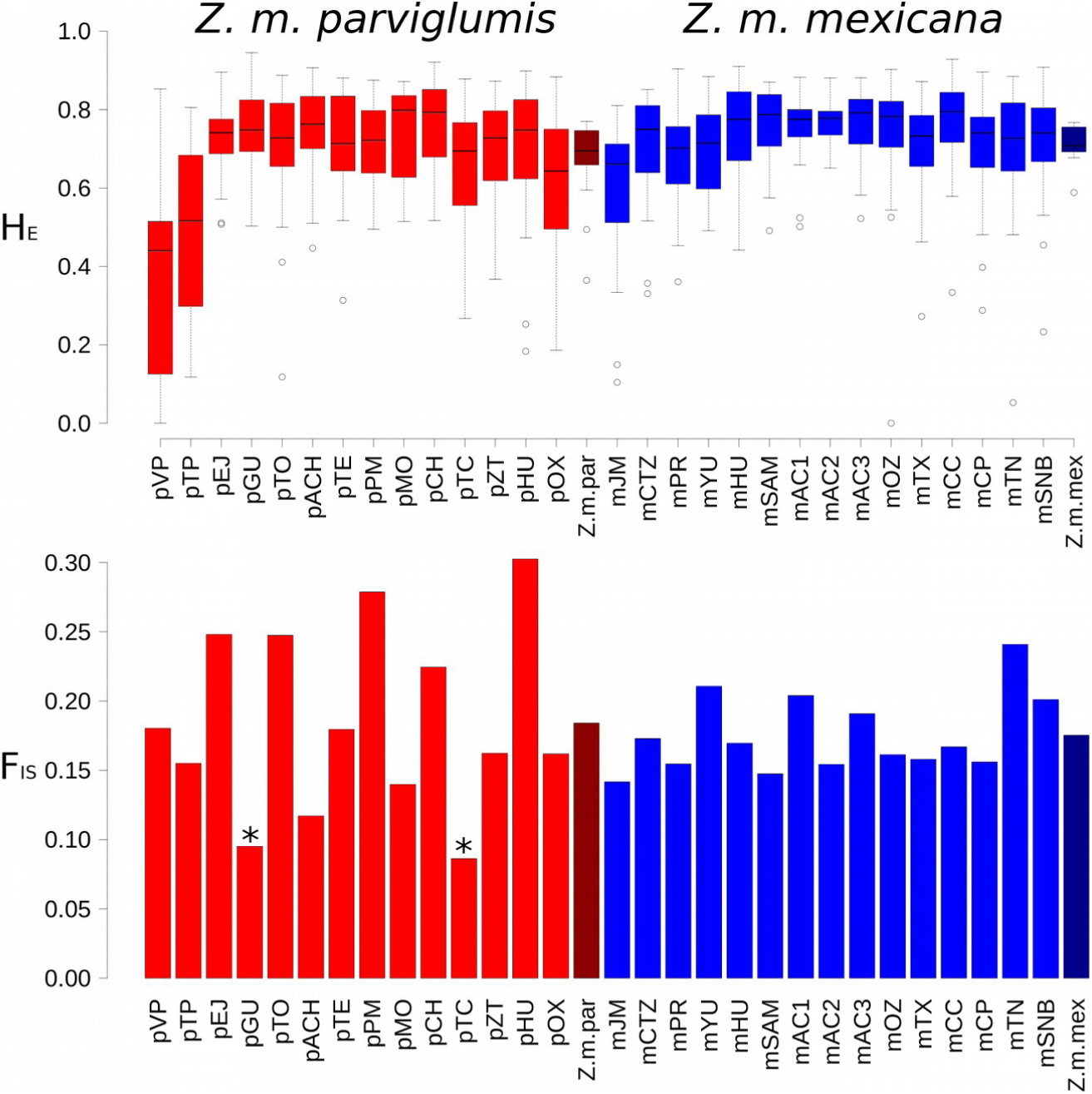
a) Expected heterozygosity per locality for *Zea mays parviglumis* (red), and *Zea mays mexicana* (blue). Z.m.par and Z.m.mex corresponds to the average value for each subespecies. b) *F*_*IS*_ values per locality, asterisks denote not-significant *F*_*IS*_ values at p>0.05. Z.m.par and Z.m.mex corresponds to the average value for each subespecies.

We found an average *F*_*IS*_ value of 0.1796, while *Z. m. parviglumis* displayed *F*_*IS*_ = 0.3374 and *Z. m. mexicana F*_*IS*_= 0.2621 (Fig. 2b, Table S4). In general, we found differences in *F*_*IS*_ values among localities (Fig. 2b), ranging from *F*_*IS*_= 0.0863 (pTC) to *F*_*IS*_= 0.3025 (pHU), with a higher fluctuation for *Z. m. parviglumis* than for *Z. m. mexicana*. Only the *Z. m. parviglumis* localities pGU and pTC were in H-W proportions. According to the *F*_*IS*_ values, Micro-Checker identified the presence of at least one null allele in each locality. However, we did not find any locus with a consistent null-allele signal across populations. Moreover, the global *F*_*ST*_ obtained with the ENA correction was not significantly different from the standard (not corrected) *F*_*ST*_ value, so we kept all loci for all posterior analyses.

Selfing estimated with RMES software showed fluctuating *sg*^*2*^ values for both teosinte subespecies (Fig. 3, Table S4); *Z. m. parviglumis* displayed higher *sg*^*2*^ values with an average *sg*^*2*^ = 0.078, and values ranging from zero (no selfing) to 0.309 per locality, while *Z. m. parviglumis* displayed an average *sg*^*2*^ = 0.052 with a range from zero to 0.133. In both subespecies, only one third of selfing estimates were significant. Also, we did not observe a spatial pattern of localities having significant selfing values.

**Figure 3.**
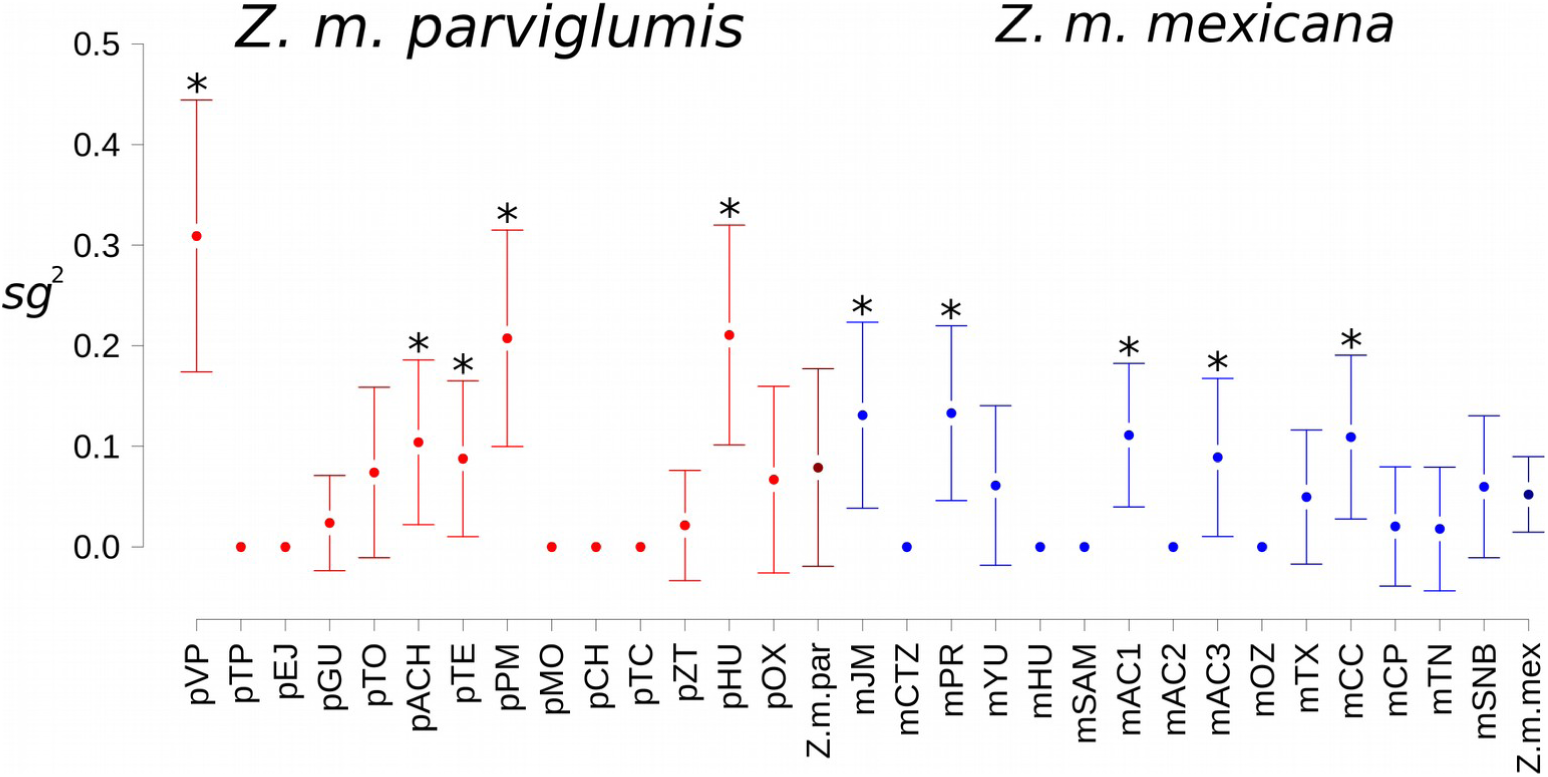
Selfing rates (*sg*^*2*^) estimated by RMES for each teosinte locality. Z.m.par and Z.m.mex corresponds to the average value for each subespecies. Bars represent standard deviation, asterisks denote values significantly different from zero.

We obtained an overall *D*_*EST*_ value of *D*_*EST*_= 0.147 between subspecies, however we found contrasting values of genetic structure among teosinte localities (Fig. 4). We observed the highest values between the *Z. m. parviglumis* pHU and pVP (*D*_*EST*_= 0.796), and the lowest values among the Michoacán localities (mAC1, mAC2, mAC3, mSAM and mHU30; mean *D*_*EST*_=0.136) of *Z. m. mexicana*. Notably, the Western-Jalisco localities (pVP and pTP) along with Oaxaca (pOX) were the most differentiated from the rest of the localities. Moreover, mJM was placed among the *Z. m. parviglumis* localities.

**Figure 4.**
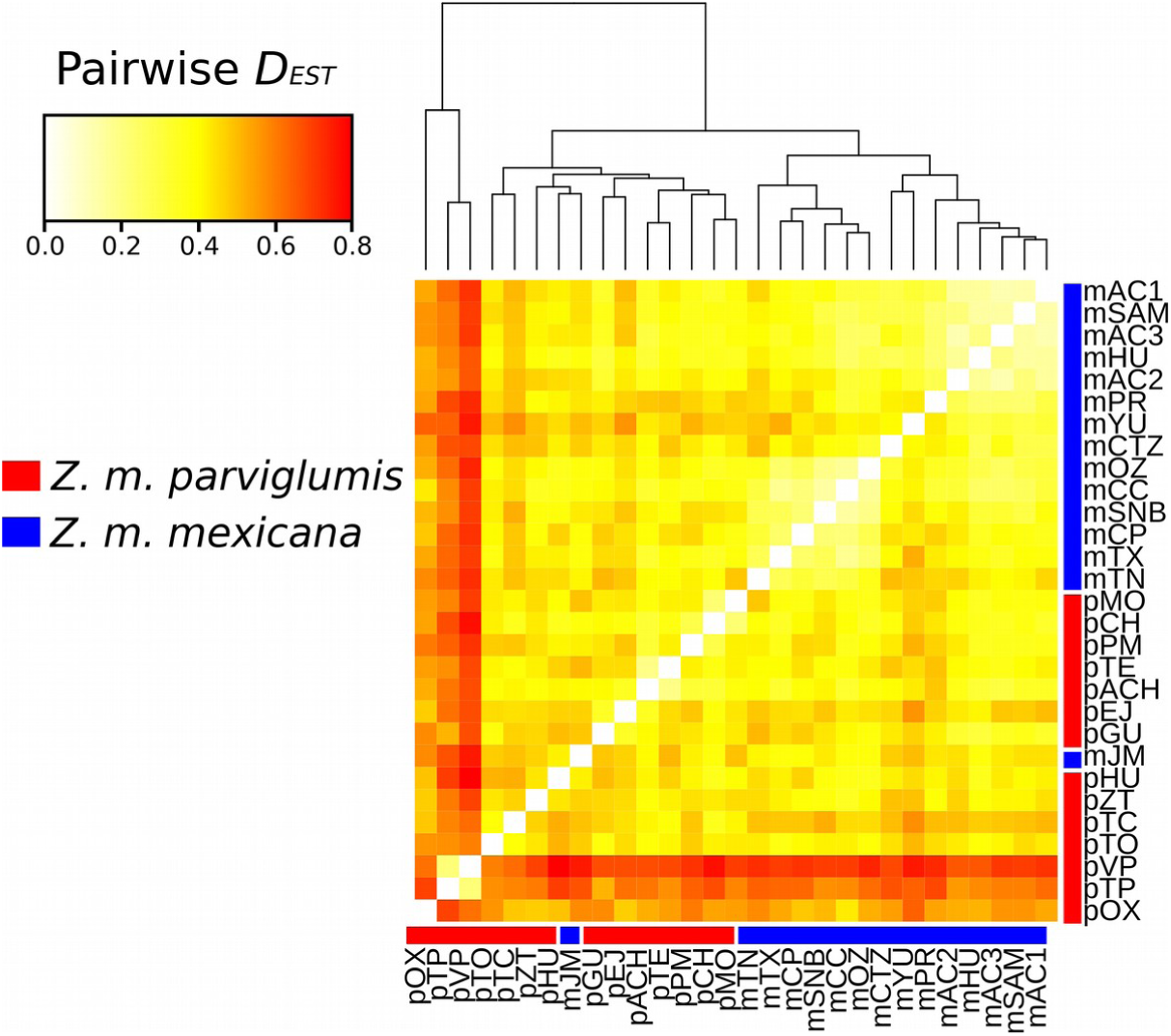
Pairwise *D*_*EST*_ values between localities. Localities were clustered using Euclidean distance with complete linkage method. The temperature color represent values from zero (white) to 0.8 (red).

The Structure analysis indicated the presence of four genetic groups (Fig. 5a). The analysis divided the localities in subspecies and each subspecies in two groups. *Z. m. parviglumis* displayed a subdivision between the Western-Jalisco localities (pVP and pTP) and the rest of the localities, while in *Z. m. mexicana*, the subdivision was between the localities of Michoacán and the rest of *Z. m. mexicana* localities. Also, the analysis showed admixed individuals between subspecies (Fig. 5a). It is worth noting that we obtained similar results when analyzing each subspecies separately (data not shown). The recent-migration analysis conducted with BayesASS between the Structure clusters yielded low migration rates (Fig. S3), and the highest rates (0.009) were obtained between Western-Jalisco and *Z. m. parviglumis* clusters.

**Figure 5.**
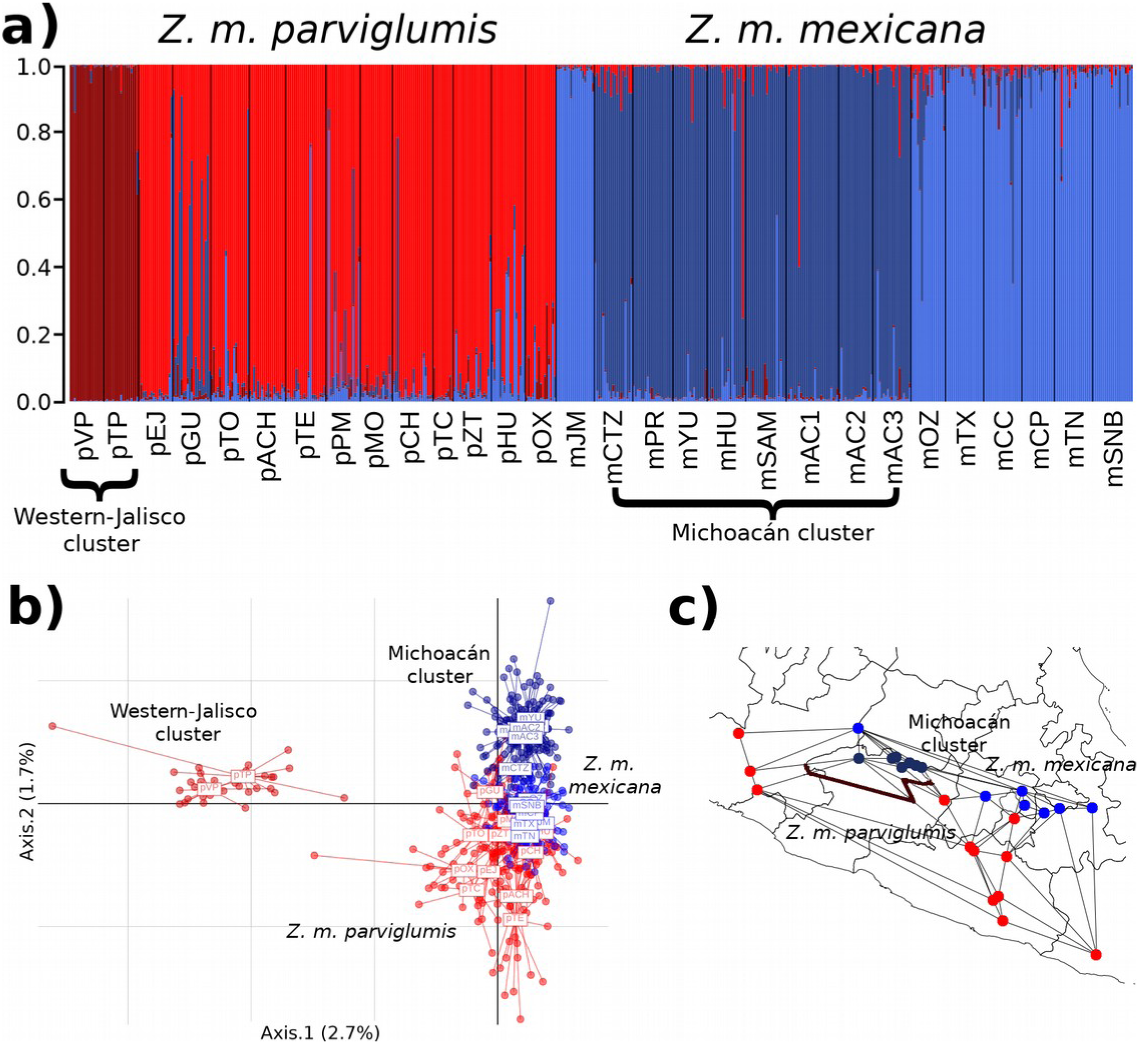
a) Bayesian assignment plot obtained with Structure for K=4. Localities were ordered from left to right according to their latitude. b) Principal Component Analysis of individuals using localities as grouping factor, dots represent individuals. c) Monmonier analysis among localities, excluding the Western-Jalisco localities (pTP and pVp, see text).

The PCA displayed the Western-Jalisco localities (pVP and pTP) as the most genetically divergent individuals (Fig. 5b), in accordance to *D*_*EST*_ and Structure results. In addition, the PCA showed genetic closeness among individuals of both subspecies, in particular from the eastern localities. On the other hand, Monmonier analysis revealed significant boundaries between the Western-Jalisco and the rest of the localities (data not shown). However, due to the possible bias generated by these genetic divergent localities (as displayed by *D*_*EST*_ and PCA), we performed the Monmonier analysis again excluding pVP and pTP, and we found significant boundaries among the Michoacán and the neighbor *Z. m. parviglumis* localities (Fig. 5c); moreover, the analysis did not detect significant boundaries within subspecies, nor between subspecies among eastern localities.

The Mantel tests between genetic distance and geographic and ecological distances displayed significant and positive correlations for both distance matrices in both subspecies (*r*>0.34, *p*<0.05, Fig. S1). Nevertheless, the Sunder analysis showed a higher probability for the environmental (*E*) model than for the geographic (*G*) and the geographic+environment (*G+E*) models in both subespecies, indicating that environmental conditions alone have a stronger influence on the change of allelic frequencies.

The VarEff tests displayed demographic changes in the four Structure clusters (Fig. 6 and Table 1). However, these analyses showed some differences in the trends and values of effective population size (*N*_*e*_) among clusters. For *Z. m. parviglumis*, the Western-Jalisco cluster showed a constant past population size, followed by steady growth, and according to its low genetic diversity, this cluster had the lowest *N*_*e*_ values (Table 1). The rest of the localities of *Z. m. parviglumis* displayed a past steady *N*_*e*_ followed by ancient demographic contraction and a posterior demographic growth (Fig. 6). In the case of *Z. m. mexicana*, the Michoacán cluster displayed a past constant *N*_*e*_, followed by the recent contraction observed for all clusters; moreover, the rest of the localities of *Z. m. mexicana* had a low constant *N*_*e*_ followed by demographic growth.

**Table 1.**
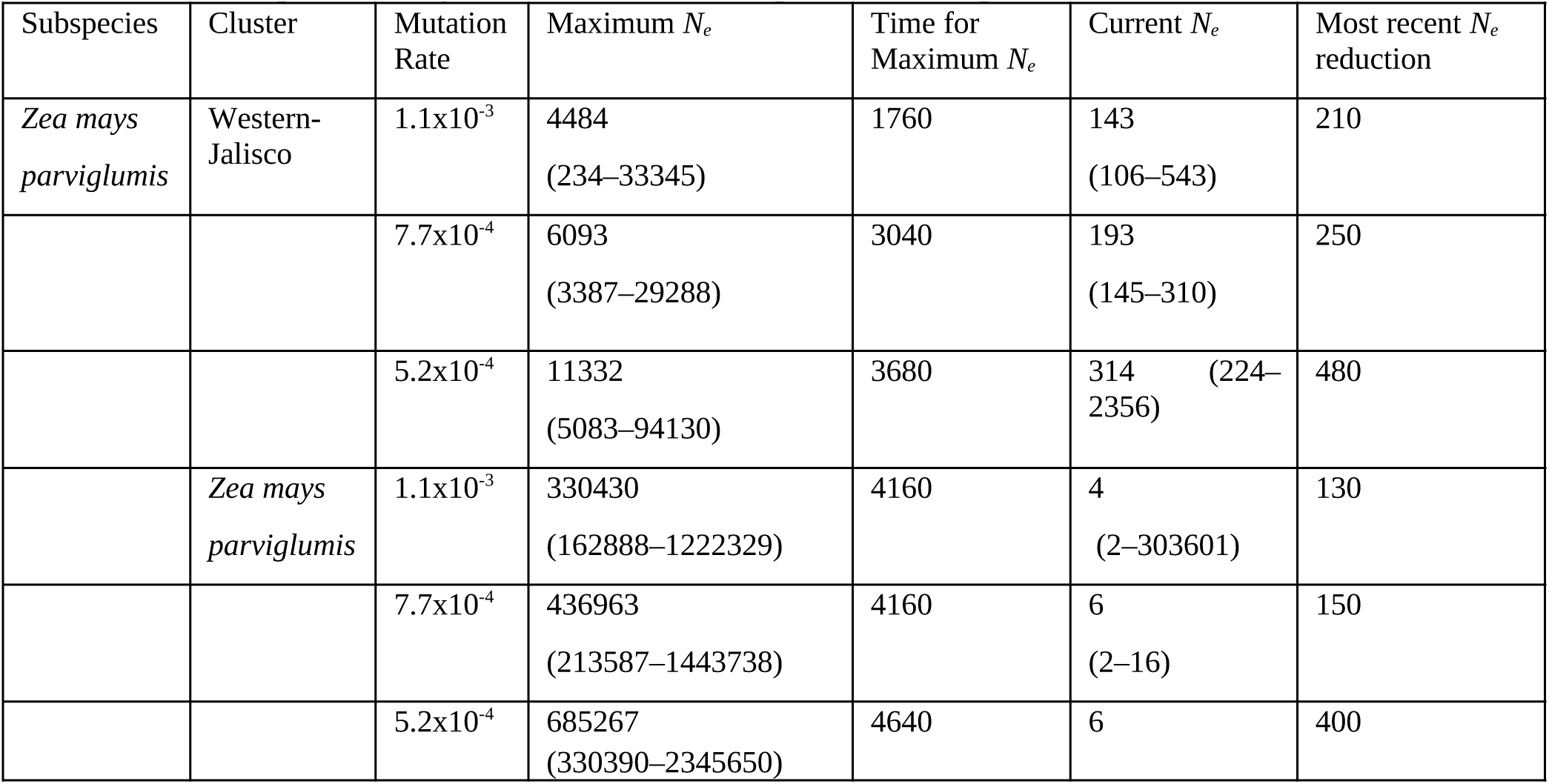
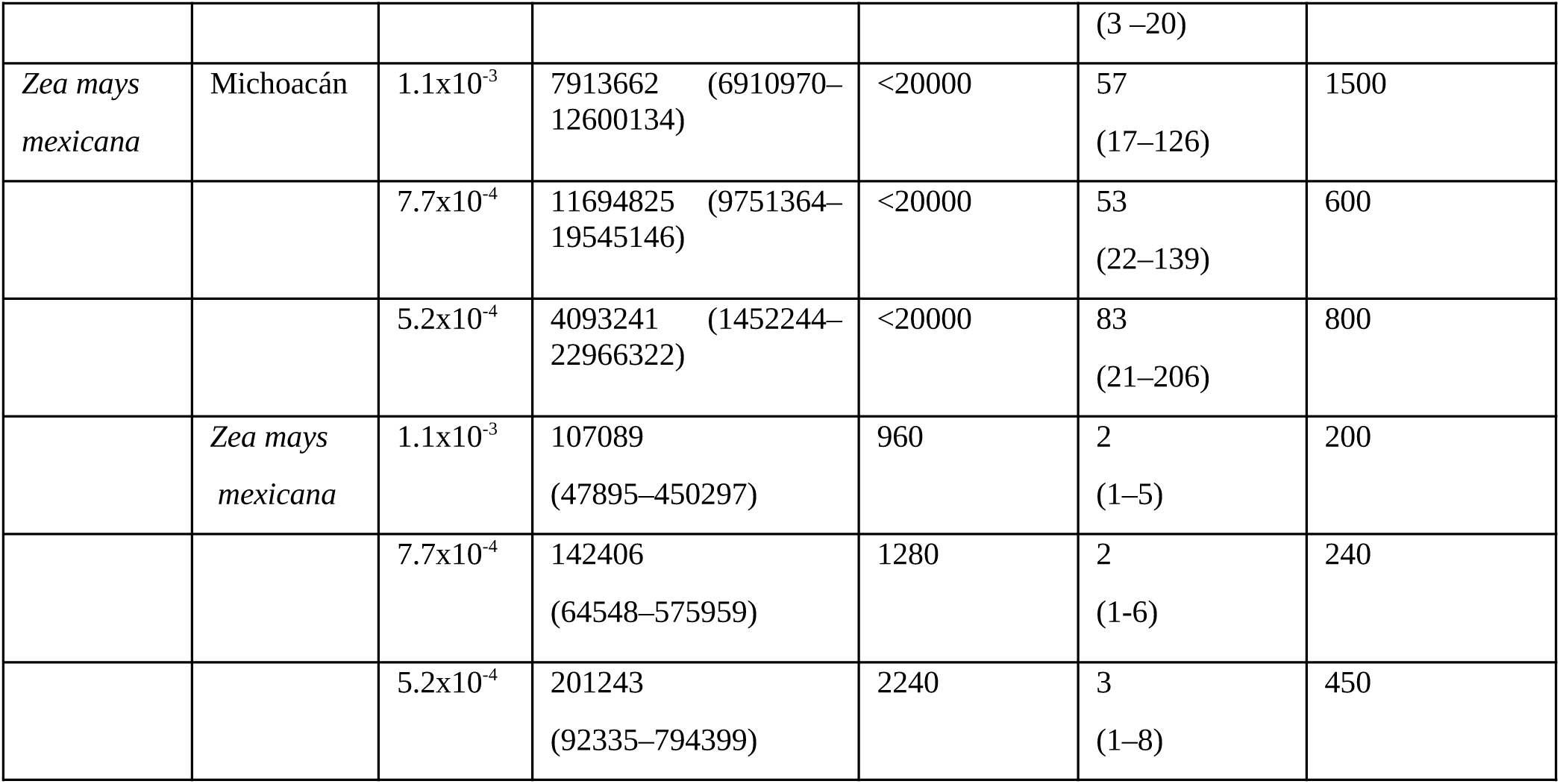
Effective population sizes (*N*_*e*_) and time of change estimated using VarEff for the clusters obtained with Structure and the range of mutation rates reported by Vigouroux et al. (2012). Time is expressed in generations. Values in parenthesis represent the 95% HPD intervals.

**Figure 6.**
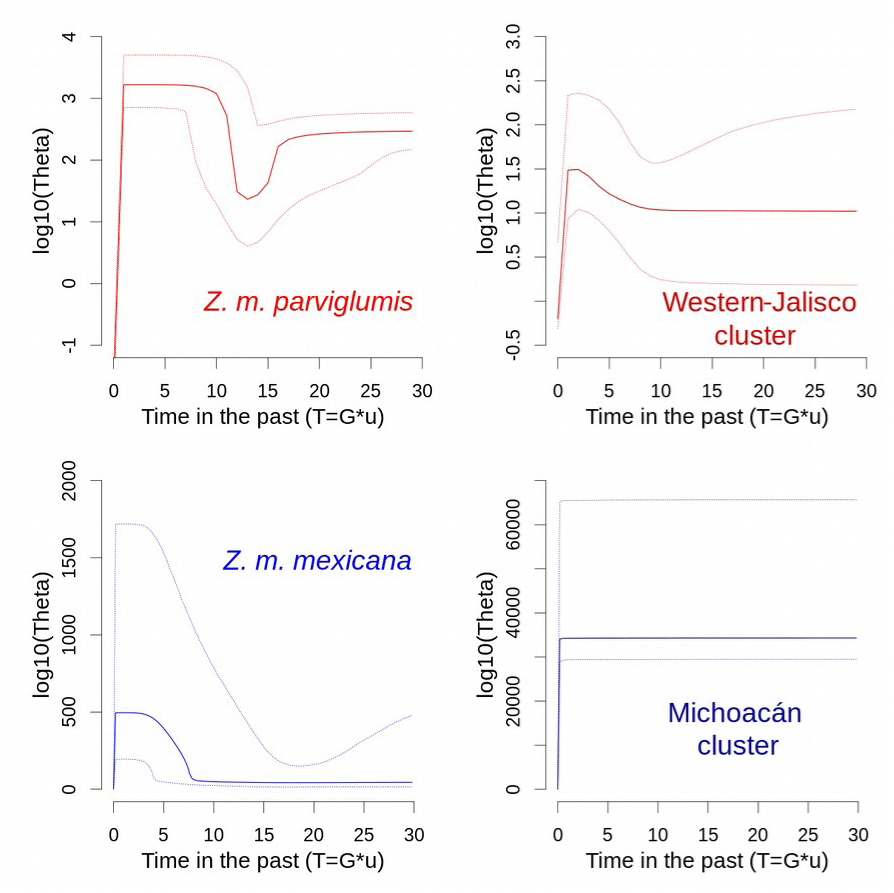
Changes in effective population sizes (*N*_*e*_) in the past estimated with VarEff for the clusters obtained with Structure. The x-axis represents time as the product of generation time per mutation rate. The y-axis represents the effective size scaled as the logarithm base 10 (log10) of Theta (4Ne*u).

Despite the differences in the demographic trends among clusters, we observed a recent and drastic *N*_*e*_ contraction 130-1,500 generations ago, resulting in a very reduced current effective size in all clusters (Table 1). Moreover, except for the Michoacán cluster, teosinte populations exhibited demographic growth after the Mid-Holocene (∼6,000 years ago) and, as displayed in Table 1, the estimated *N*_*e*_ suggested large population sizes in the past (as large as <11×10^6^ for the Michoacán cluster).

SDM displayed allopatric current potential distribution areas among subspecies (Fig. 7), showing only ∼3.5% overlap in the boundaries of each subspecies distribution. Furthermore, both subspecies exhibited changes in their distribution areas across four different time periods. In the LIG (∼120kys) period we estimated a small distribution area, while for the LGM (∼22kys) we observed an increase in the potential distribution area in both subspecies. Moreover, for the MID (∼6kys) we observed an acute reduction in the potential distribution, followed by an increment to the present (Fig. 7). The reduction in the predicted distribution area for the MID may be explained by the contrasting environmental conditions at this period (Fig. S2).

**Figure 7.**
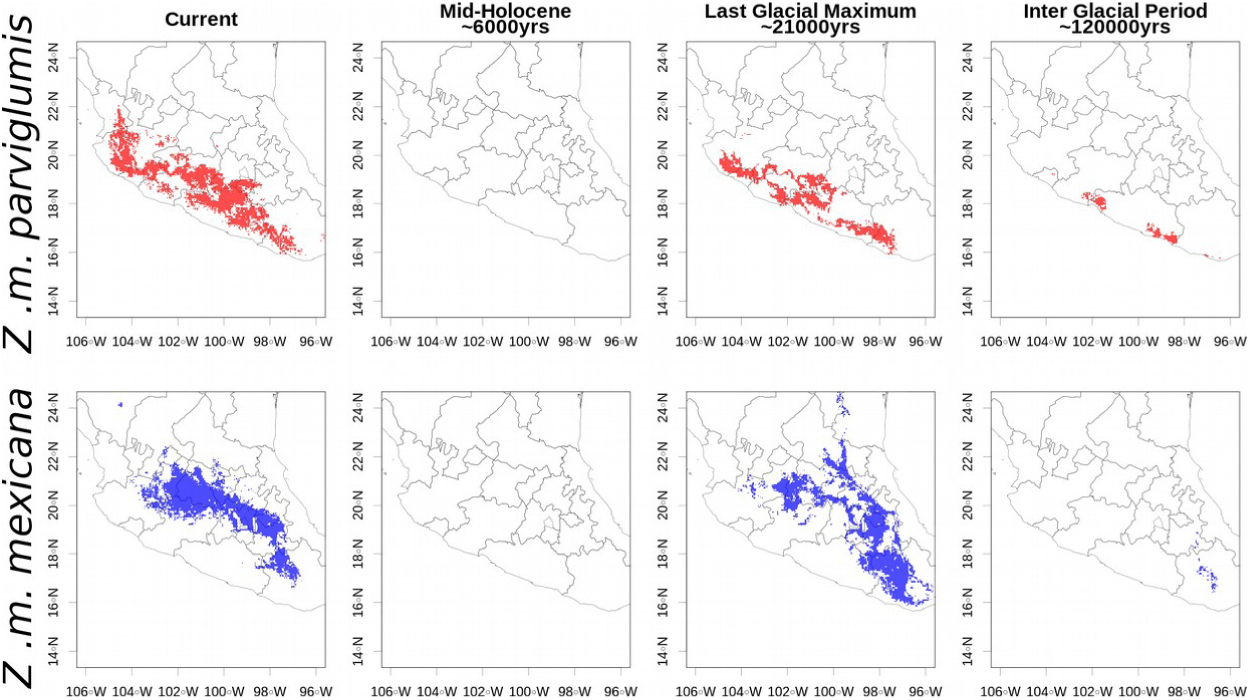
Species Distribution Ensemble Models for *Zea mays parviglumis* (red) and *Zea mays mexicana* (blue) considering current conditions, mid-holocene (MID 6,000 years), the last glacial maximum (LGM 21,000 years) and last interglacial period (LIG 120,000 years).

## Discussion

Our analyses in current populations of *Z. m. parviglumis* and *Z. m. mexicana* revealed that local factors like selfing rates, isolation and historical processes as climate change, have shaped the levels and distribution of their genetic diversity. At the local scale, the number of alleles and heterozygosity were consistent with previous reports on teosinte (Matsuoka *et al*., 2002; Fukunaga *et al*., 2005). Nevertheless, the low diversity detected in the Western-Jalisco cluster could be related to genetic drift due to genetic isolation, as this was confirmed by the low *N*_*e*_ estimated for these localities (Table 1). Besides, *F*_*IS*_ values showed that most teosinte localities had from moderate to high levels of heterozygote deficiency and only two *Z. m. parviglumis* localities were in H-W proportions, suggesting ongoing evolutionary process like genetic drift and inbreeding affecting allele frequencies. Even though positive *F*_*IS*_ values may indicate within-localities inbreeding (Wright, 1965), the generalized heterozygote deficiency signal that we recovered, could also be caused by Wahlund effect (excess of observed homozygotes resulting from sampling a structured population) produced by local genetic structure. Moreover, the RMES analysis showed no correlation between selfing and *F*_*IS*_ values and we found that neighbouring localities did not have similar selfing values. For instance, localities from the Michoacán cluster had the lowest *D*_*EST*_ values (suggesting some degree of genetic connectivity) while their selfing values s*g*^*2*^ ranged from zero to 0.233 (Fig. 3). A previous study reported high levels of outcrossing at local scales (Van Heerwaarden *et al*., 2010); however, the authors also suggested that the fine-scale environmental features, like topography and wind, may influence pollen and seed dispersal generating local genetic structure. The lack of spatial arrangement of *F*_*IS*_ and s*g*^*2*^, could be associated with limited seed dispersion that promote local selfing and genetic structure. In this sense, the teosinte is considered as a Pleistocene species adapted to seed dispersal by large herbivores, and although livestock grazes on their plants in the present, their range of dispersion is limited by the human agricultural niche construction (mainly land crops) (Webster, 2011), restricting dispersion to a short distances. This, in turn, leads to the foundation of small and fragmented populations genetically isolated.

On a global scale, we found relatively high levels of genetic structure, and accordingly, low estimates of gene flow. Furthermore, we observed that the Oaxaca locality (pOX) and Western-Jalisco cluster (pVP and pTP) were the most differentiated localities. While this was expected for pOX (as this locality was the most geographically distant), the Western-Jalisco cluster was highly differentiated even in the presence of relatively close localities (for example, pGU, pEJ, and pTO <80 km; Fig. 1 and 4). Moreover, the mJM locality is situated in a neighboring area near the Michoacán cluster localities (Fig. 5); yet, it was highly structured from the other *Z. m mexicana* localities. Our results suggest that spatial distribution alone is not the determinant factor for the geographical distribution of genetic diversity on teosinte populations. In particular, the Sunder analysis showed that the environment provides a better explanation for values of genetic differentiation in teosinte than geographic distance. This result supports the signatures of local adaptation detected by Aguirre-Liguori *et al*., (2019) using a large SNPs data set. Hence, our results indicate that adaptation to local conditions is promoting the genetic divergence among teosinte populations and subspecies, despite the presence of dispersal and gene flow. This hypothesis is supported by the differences in the environmental conditions encountered in the distribution of the genetic clusters (Fig. S4), and the reduced proportion of suitable areas shared by subspecies, as shown by current SDM projections (Fig. 7). Another factor that may reduce the homogenization of genetic diversity via gene flow is the variable outcrossing rates observed that preclude admixture with external populations. Moreover, we observed that our genetic clusters of *Z. m. mexicana* correspond to the “geographic races” Chalco (main *Z. m. mexicana* cluster) and Central Plateau (Michoacán cluster) proposed by Gonzalez et al. (2018) based exclusively on environmental features.

On the other hand, the drastic reduction in the predicted area during MID may be produced by the marked differences on seven of the ten bioclimatic variables in that period, (mainly rain regimes, Fig. S2). Furthermore, the observed changes in the historical effective population sizes (*N*_*e*_) concurred with SDM analyses. For three of the four genetic clusters, we found a demographic increase with a peak of the *N*_*e*_ posterior to MID (Fig. 6), indicating that demographic growth may have been promoted by a period of suitable conditions after a period of non-favorable conditions. Although we observed a fluctuation of suitable distribution areas across the four analyzed periods (a very reduced area in LIG, an increase in the suitable area in the LGM, a drastic reduction in MID, and a new increase in the current-time), microsatellite markers showed resolution for the last ∼20,000 generations, thus limiting the analyses to more recent events than the LGM. Interestingly, the Michoacán cluster displayed an ancient constant *N*_*e*_, suggesting a negligible impact from the MID environmental shift. This trend could be related to the possible existence of a post-glacial refugium (not detected by our projections) that, coupled with the genetic connectivity observed in this area, allowed large *N*_*e*_ across time. Another possible explanation is related to the particular environmental requirements of this cluster (Fig. S4). If that is the case, projections performed in the complete database of *Z. m. mexicana* may result in a biased distribution modeling of the Michoacán cluster. To evaluate this hypothesis, however, it will be necessary to perform extensive sampling and posterior assignment analysis to collect enough georeferenced data and to obtain reliable projections.

Finally, the low *N*_*e*_ values observed as a consequence of a recent demographic contraction should be interpreted with caution. The severe bottleneck observed may be the outcome of the exploitation of teosinte as a food source in the pre-Columbian Mesoamerican populations (Webster, 2011), and more recently, the result of displacement of teosinte due to maize production. However, in some of our estimations, the effective size contraction continued as recently as ten generations ago, resulting in an *N*_*e*_ of almost zero. This is likely a result of an inaccurate adjustment of *N*_*e*_ caused by a complex demographic history and a very recent and severe demographic contraction, causing the algorithm not to achieve an accurate estimation in very recent time scale.

## Conclusions

Our results showed that historical and contemporary factors at local and global scale shaped the genetic diversity of teosinte. While local and contemporary factors cause differences in mating patterns and genetic structure, the historical factors shaped the amount and distribution of genetic diversity of teosinte populations. In this sense we remark the importance of including data from different temporal and spatial scales in order to obtain a better picture of the mechanisms that affect genetic diversity. Also, we found evidence that suggests that the genetic lineages found could be the result of local adaptation as previously suggested by other authors (Aguirre-Liguori *et al.*, 2017; Fustier *et al.*, 2017; Aguirre-Liguori *et al.*, 2019). This adaptability to specific environmental and agroecological conditions permits the wide ecological conditions inhabited by teosinte.

Even though microsatellites are considered neutral markers, we recovered strong signals of genetic structure that suggest both an ongoing process of genetic drift, and local adaptation to environmental conditions that maintain different genetic lineages despite possible introgression. We also found that teosinte populations have undergone multiple genetic bottlenecks, and that, along with the detected genetic structure, make teosinte populations prone to local extinction. As a consequence, we may be losing an important genetic reservoir for the permanence of teosinte, but also genetic variants that could be relevant to maize improvement and adaptation to global environmental change.

## Acknowledgments

The original sampling of the populations and of the overall study was designed with the help and enthusiasm of Drs. Brandon S. Gaut, Maud I. Tenaillon, Valeria Souza and Salvador Montes-Hernández. We thank L. Espinosa-Asuar and M. Rosas-Barrera for their technical assistance, Alberto Villasante Barahona for his help in generating the microsatellite datasets and Silvia Barrientos for her support during this investigation. We also thank Dr. Gabriela Castellanos-Morales for her valuable comments for the improvement of the manuscript. This work was funded by grants CB2011/167826 (CONACYT Investigación Científica Básica), CN 10 393 (UC MEXUS CONACYT), and M12 A03 ECOS Nord France CONACYT-ANUIES.

## Supplementary Materials

### Supplemental Figures

**Figure S1.**
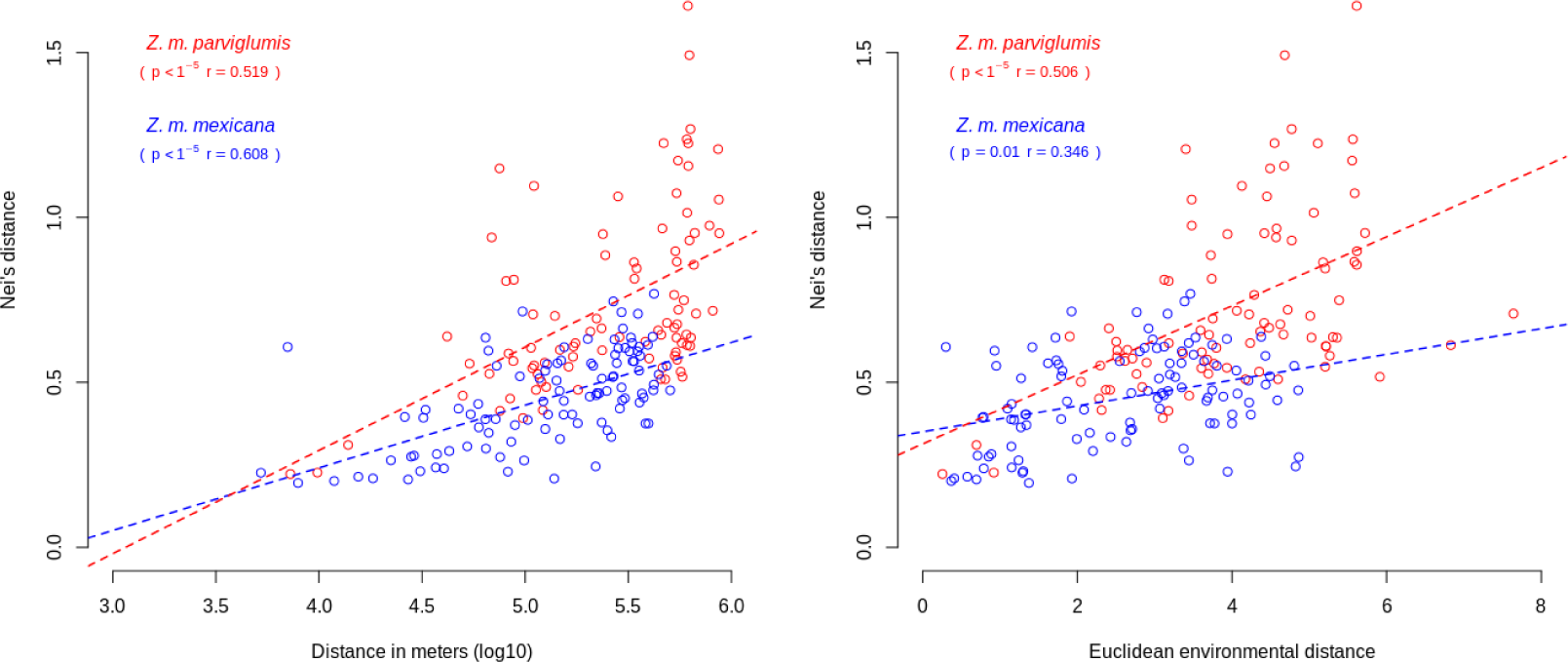
Mantels tests for genetic distance matrix against geographic distance, and environmental distance for *Zea mays parviglumis* (in red) and *Zea mays mexicana* (in blue).

Figure S2. Boxplots of the environmental variables included for the SDM projections for four time periods (current, MID, LGM and LIG).

Figure S3. Migration rates among Structure clusters calculated with BayesASS software.

**Figure S4.**
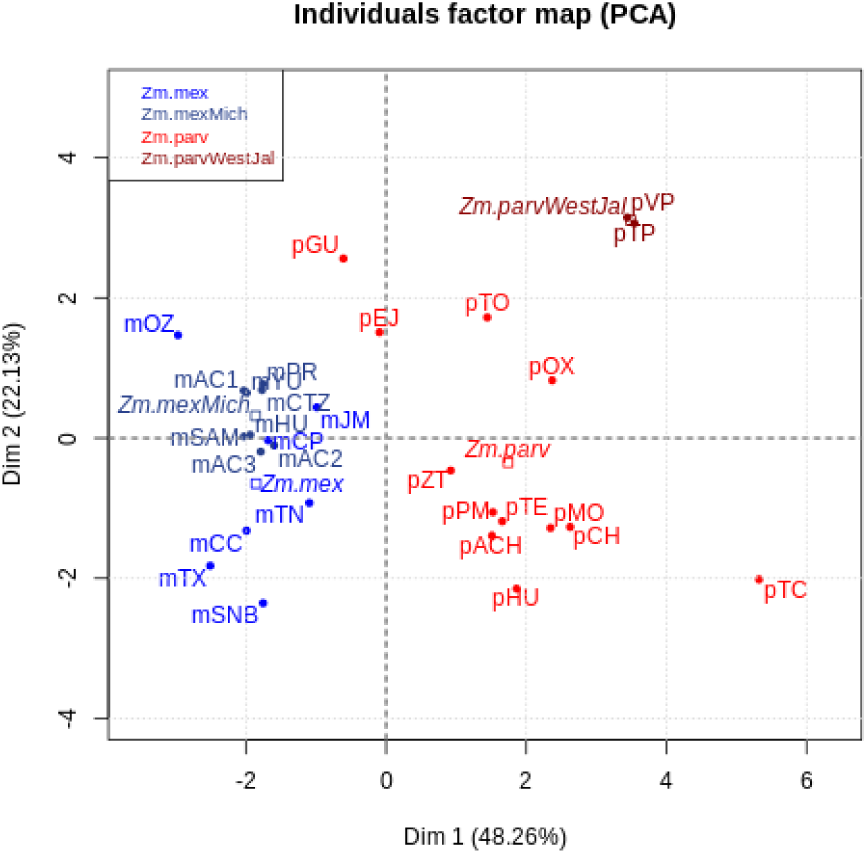
Principal Components Analysis of the bioclim variables for teosinte localities analyzed in this study.

## Supplemental tables

**Table S1.**
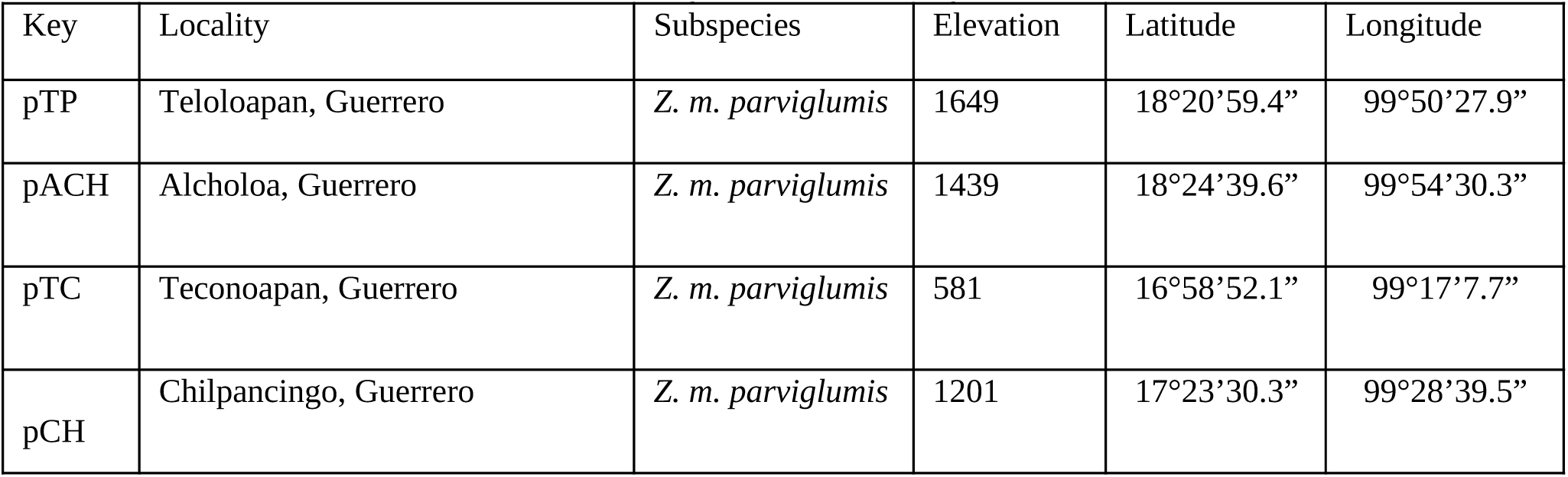
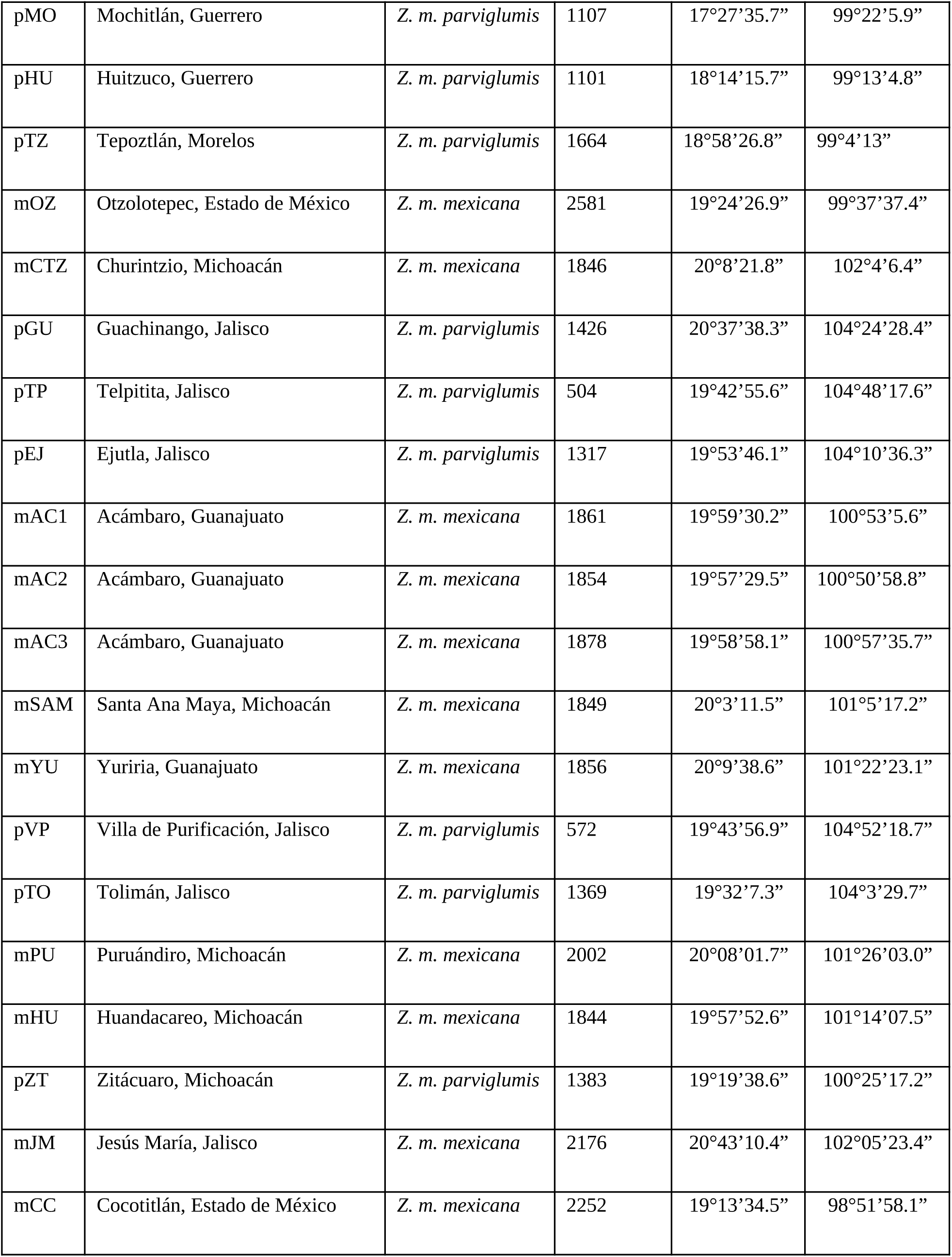
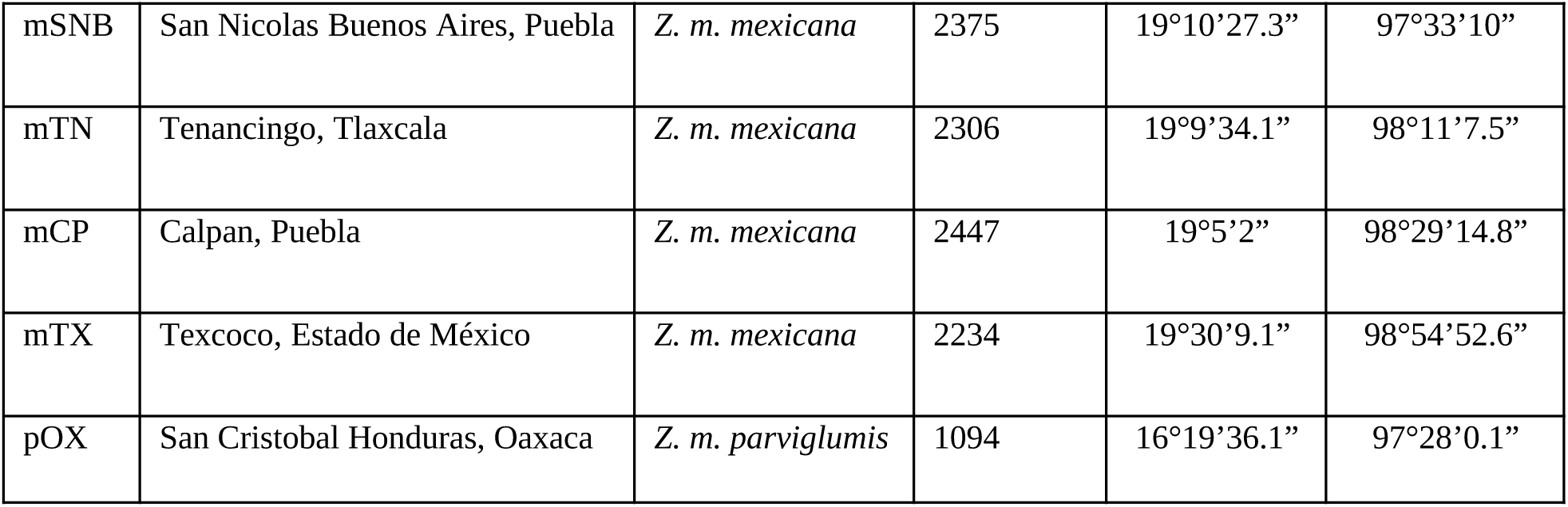
Information of the localities analyzed in this study.

**Table S2.**
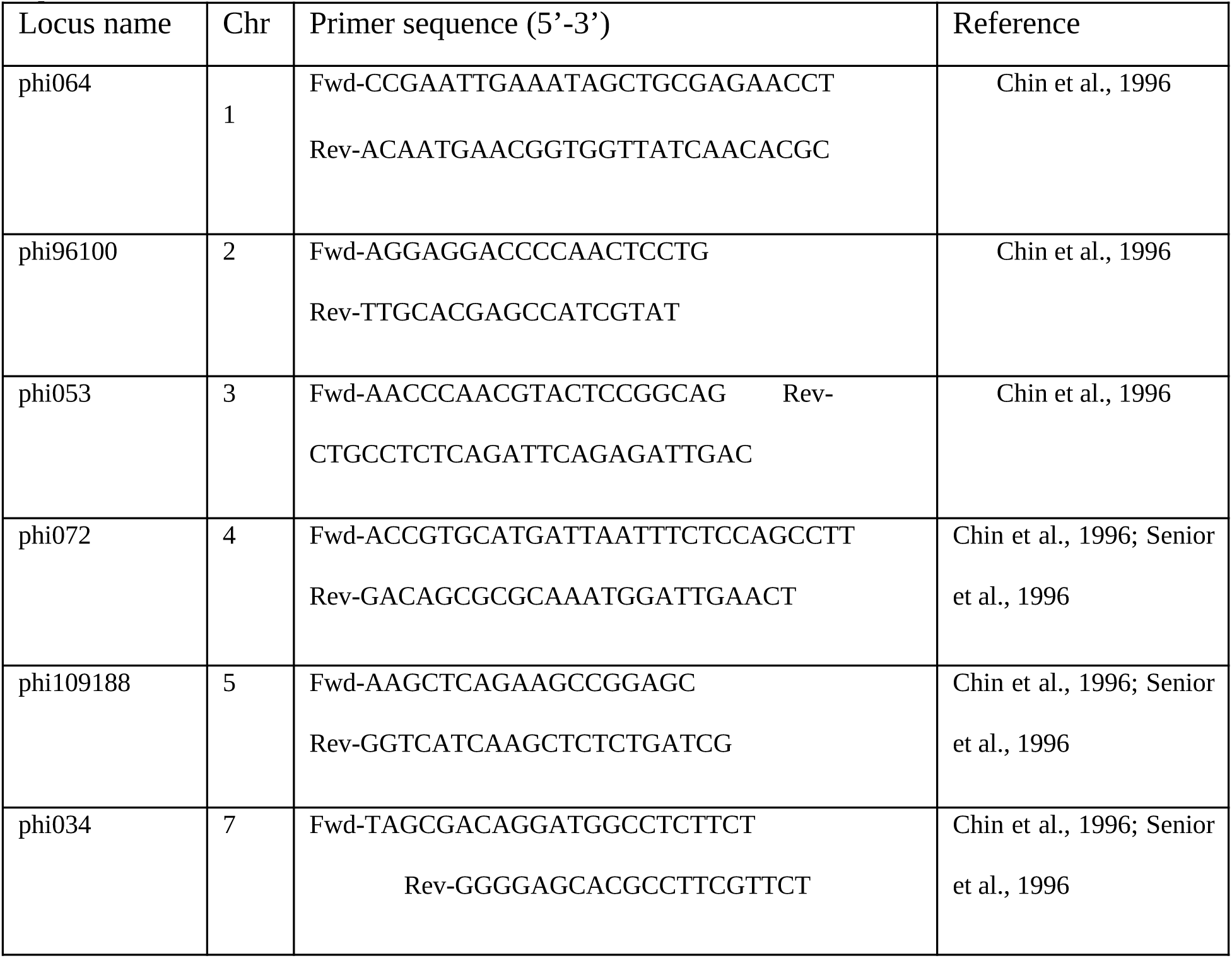
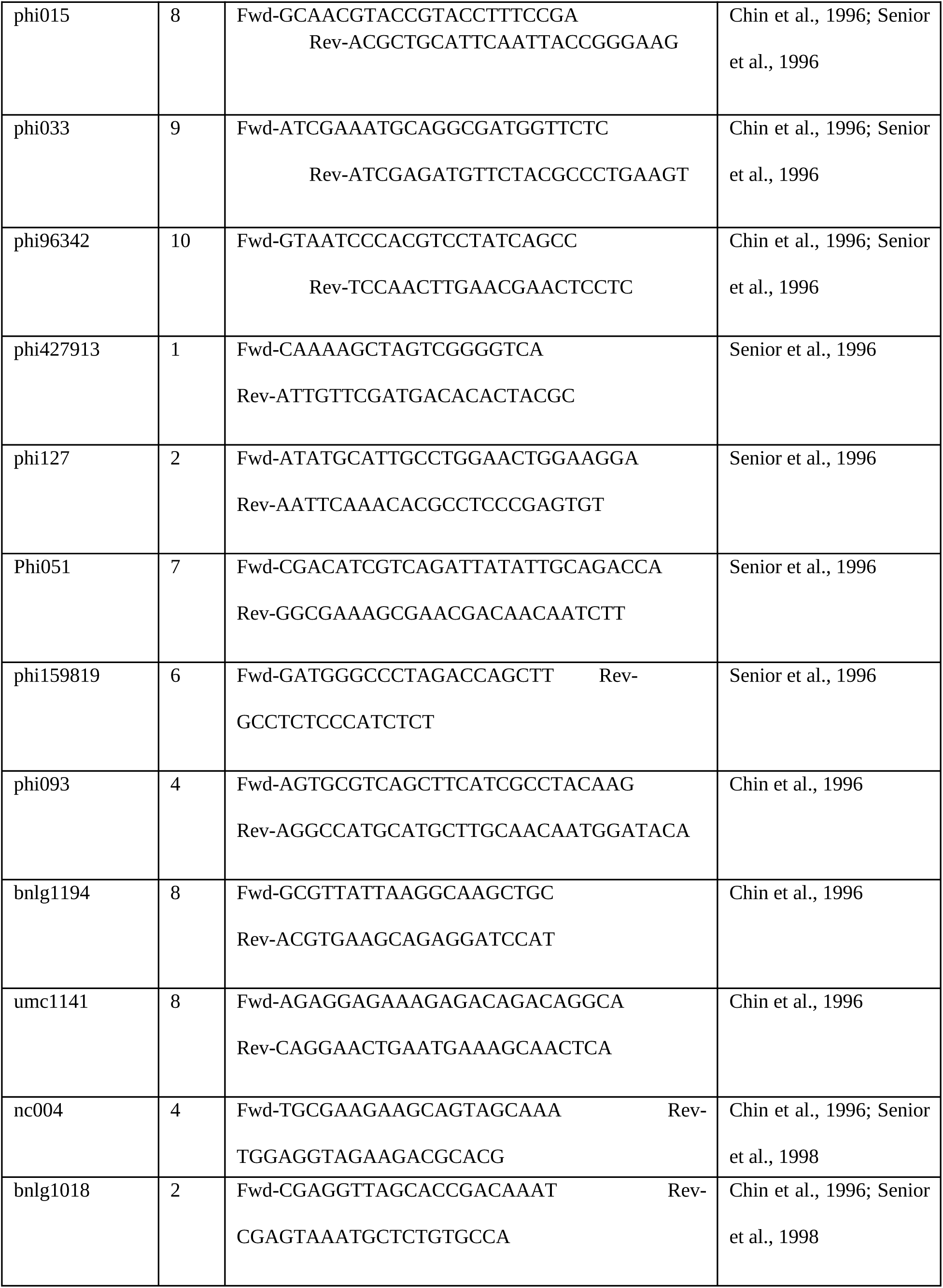
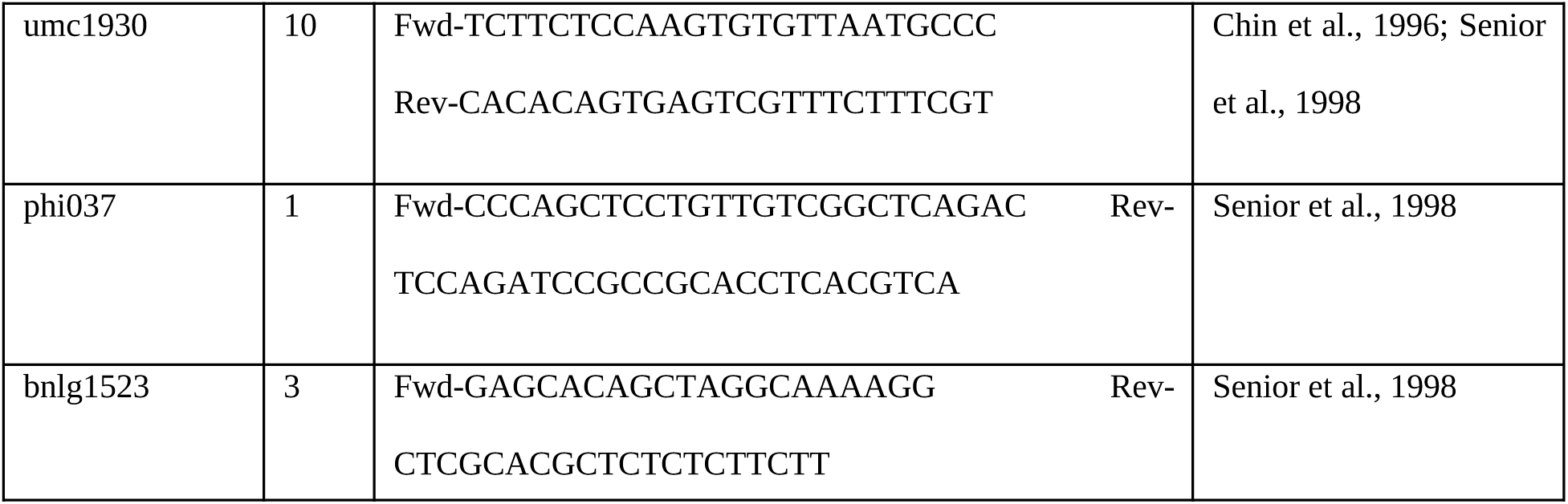
Locus name for the 22 microsatellite loci used in this study, chromosomal position (Chr) in *Zea mays* according to reported on MaizeGDB (http://www.maizegdb.org/), primer sequences, and reference.

**Table S4.** Values of expected heterozygosity (*H*_*E*_), observed heterozygosity (*H*_*O*_), fixation index (*F*_*IS*)_ and selfing (*sg*^*2*^) for each teosinte locality.

| Locality | $H_E$ | $H_O$ | $F_{IS}$ | $sg^2$ |
| --- | --- | --- | --- | --- |
| pVP | 0.36435 | 0.28632 | 0.1803 | 0.30904 |
| pTP | 0.49409 | 0.41591 | 0.1551 | 0 |
| pEJ | 0.72746 | 0.54663 | 0.2481 | 0 |
| pGU | 0.74926 | 0.66433 | 0.0951 | 0.02385 |
| pTO | 0.68631 | 0.50762 | 0.2475 | 0.07412 |
| pACH | 0.74871 | 0.65616 | 0.117 | 0.104 |
| pTE | 0.71214 | 0.58397 | 0.1795 | 0.08773 |
| pPM | 0.7025 | 0.50028 | 0.2787 | 0.20735 |
| pMO | 0.74663 | 0.64286 | 0.1398 | 0 |
| pCH | 0.76996 | 0.59383 | 0.2244 | 0 |
| pTC | 0.65878 | 0.60253 | 0.0863 | 0 |
| pZT | 0.68678 | 0.57298 | 0.1622 | 0.02147 |
| pHU | 0.6851 | 0.47605 | 0.3025 | 0.2106 |
| pOX | 0.59499 | 0.49343 | 0.1618 | 0.06703 |
| mJM | 0.58855 | 0.50467 | 0.1417 | 0.13099 |
| mCTZ | 0.69755 | 0.5758 | 0.1729 | 0 |
| mPR | 0.67688 | 0.57091 | 0.1547 | 0.13302 |
| mYU | 0.70021 | 0.5513 | 0.2106 | 0.06118 |
| mHU | 0.75793 | 0.62758 | 0.1695 | 0 |
| mSAM | 0.75529 | 0.64286 | 0.1476 | 0 |
| mAC1 | 0.7532 | 0.59399 | 0.2041 | 0.1111 |
| mAC2 | 0.76713 | 0.64706 | 0.1543 | 0 |
| mAC3 | 0.75575 | 0.6043 | 0.1909 | 0.08903 |
| mOZ | 0.72711 | 0.61046 | 0.1612 | 0 |
| mTX | 0.6942 | 0.58386 | 0.158 | 0.04966 |
| mCC | 0.76526 | 0.63755 | 0.1669 | 0.10916 |
| mCP | 0.68992 | 0.56629 | 0.1561 | 0.02043 |
| mTN | 0.68381 | 0.50982 | 0.2409 | 0.01796 |
| mSNB | 0.70817 | 0.55983 | 0.2011 | 0.05991 |

