## Supplemental Figures S1-S4 for "The role of environment, local adaptation and past climate fluctuation on the amount and distribution of genetic diversity in the teosinte in Mexico"

### Supplementary Materials

#### Supplemental Figures

Figure S1

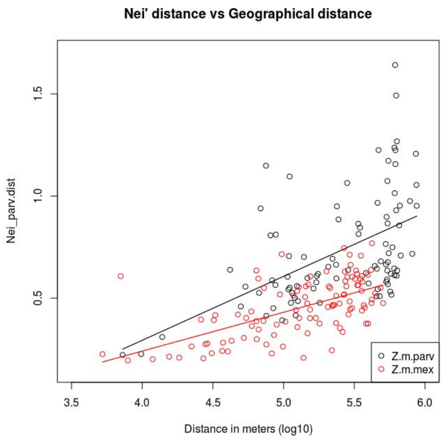


Figure S2

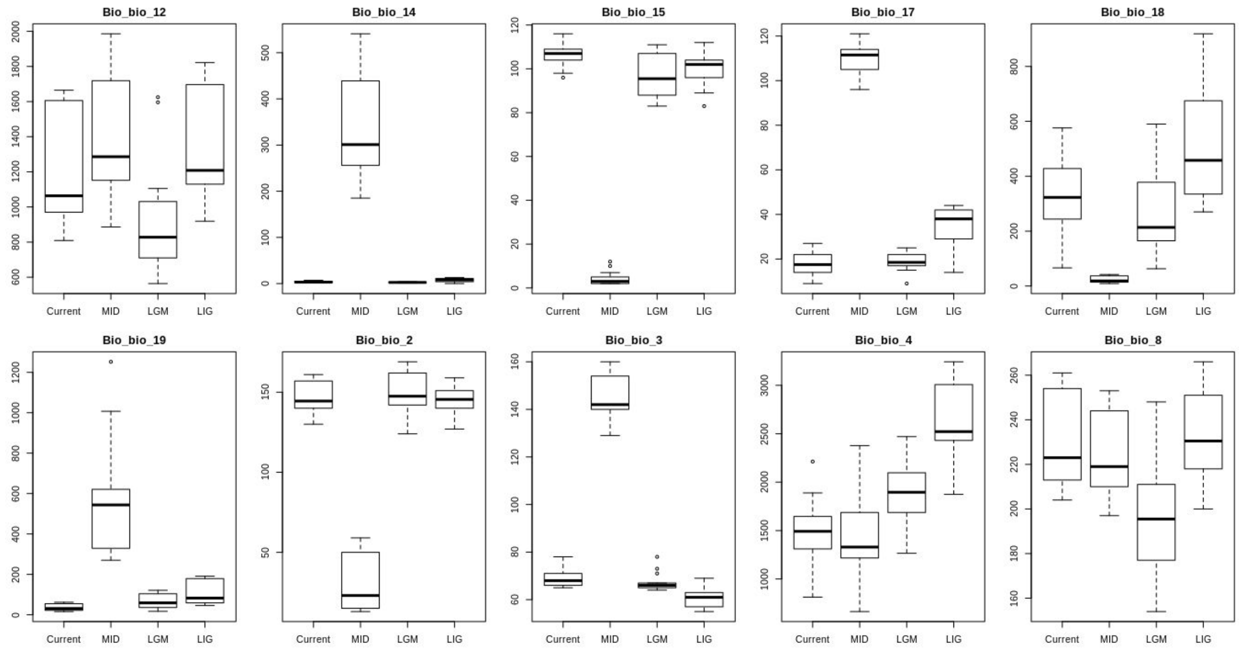


Figure S3

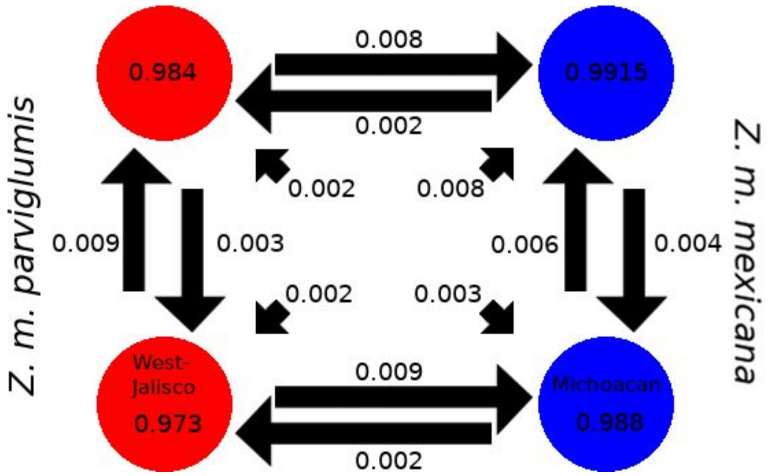


Figure S4

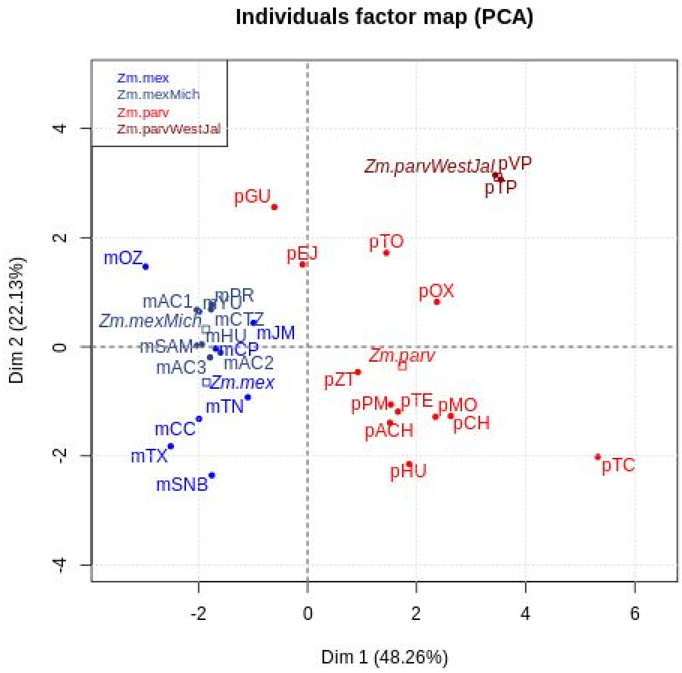


Supplemental figures captions

Figure S1. Mantels tests for genetic distance matrix against a) geographic distance, and b) environmental distance.

Figure S2. Boxplots of  the included environmental variables for the four periods of time.

Figure S3. Migration rates among Structure clusters calculated with BayesASS software.

Figure S4. Principal Components Analysis of the bioclim variables for the teosintes localities analyzed in this study.

Supplemental tables

Table S1. Information of the localities analized in this work.

| Key | Locality | Subspecies | Elevation | Latitude | Longitude |
|---|---|---|---|---|---|
| pTP | Teloloapan, Guerrero | *Z. m. parviglumis* | 1649 | 18°20’59.4” | 99°50’27.9” |
| pACH | Alcholoa, Guerrero | *Z. m. parviglumis* | 1439 | 18°24’39.6” | 99°54’30.3” |
| pTC | Teconoapan, Guerrero | *Z. m. parviglumis* | 581 | 16°58’52.1” | 99°17’7.7” |
| pCH | Chilpancingo, Guerrero | *Z. m. parviglumis* | 1201 | 17°23’30.3” | 99°28’39.5” |
| pMO | Mochitlán, Guerrero | *Z. m. parviglumis* | 1107 | 17°27’35.7” | 99°22’5.9” |
| pHU | Huitzuco, Guerrero | *Z. m. parviglumis* | 1101 | 18°14’15.7” | 99°13’4.8” |
| pTZ | Tepoztlán, Morelos | *Z. m. parviglumis* | 1664 | 18°58’26.8” | 99°4’13” |
| mOZ | Otzolotepec, Estado de México | *Z. m. mexicana* | 2581 | 19°24’26.9” | 99°37’37.4” |
| mCTZ | Churintzio, Michoacán | *Z. m. mexicana* | 1846 | 20°8’21.8” | 102°4’6.4” |
| pGU | Guachinango, Jalisco | *Z. m. parviglumis* | 1426 | 20°37’38.3” | 104°24’28.4” |
| pTP | Telpitita, Jalisco | *Z. m. parviglumis* | 504 | 19°42’55.6” | 104°48’17.6” |
| pEJ | Ejutla, Jalisco | *Z. m. parviglumis* | 1317 | 19°53’46.1” | 104°10’36.3” |
| mAC1 | Acámbaro, Guanajuato | *Z. m. mexicana* | 1861 | 19°59’30.2” | 100°53’5.6” |
| mAC2 | Acámbaro, Guanajuato | *Z. m. mexicana* | 1854 | 19°57’29.5” | 100°50’58.8” |

| | | | | | |
|---|---|---|---|---|---|
| mAC3 | Acámbaro, Guanajuato | *Z. m. mexicana* | 1878 | 19°58’58.1” | 100°57’35.7” |
| mSAM | Santa Ana Maya, Michoacán | *Z. m. mexicana* | 1849 | 20°3’11.5” | 101°5’17.2” |
| mYU | Yuriria, Guanajuato | *Z. m. mexicana* | 1856 | 20°9’38.6” | 101°22’23.1” |
| pVP | Villa de Purificación, Jalisco | *Z. m. parviglumis* | 572 | 19°43’56.9” | 104°52’18.7” |
| pTO | Tolimán, Jalisco | *Z. m. parviglumis* | 1369 | 19°32’7.3” | 104°3’29.7” |
| mPU | Puruándiro, Michoacán | *Z. m. mexicana* | 2002 | 20°08’01.7” | 101°26’03.0” |
| mHU | Huandacareo, Michoacán | *Z. m. mexicana* | 1844 | 19°57’52.6” | 101°14’07.5” |
| pZT | Zitácuaro, Michoacán | *Z. m. parviglumis* | 1383 | 19°19’38.6” | 100°25’17.2” |
| mJM | Jesús María, Jalisco | *Z. m. mexicana* | 2176 | 20°43’10.4” | 102°05’23.4” |
| mCC | Cocotitlán, Estado de México | *Z. m. mexicana* | 2252 | 19°13’34.5” | 98°51’58.1” |
| mSNB | San Nicolas Buenos Aires, Puebla | *Z. m. mexicana* | 2375 | 19°10’27.3” | 97°33’10” |
| mTN | Tenancingo, Tlaxcala | *Z. m. mexicana* | 2306 | 19°9’34.1” | 98°11’7.5” |
| mCP | Calpan, Puebla | *Z. m. mexicana* | 2447 | 19°5’2” | 98°29’14.8” |
| mTX | Texcoco, Estado de México | *Z. m. mexicana* | 2234 | 19°30’9.1” | 98°54’52.6” |
| pOX | San Cristobal Honduras, Oaxaca | *Z. m. parviglumis* | 1094 | 16°19’36.1” | 97°28’0.1” |

Table S2. Microsatellite loci used in this study, chromosomal position in *Zea mays* according to reported on MaizeGDB (http://www.maizegdb.org/), primer sequences, and references.

| Locus name | Chr | Primer sequence (5’-3’) | Reference |
|---|---|---|---|

| | | | |
|---|---|---|---|
| phi064 | 1 | Fwd-CCGAATTGAAATAGCTGCGAGAACCT<br>Rev-ACAATGAACGGTGGTTATCAACACGC | Chin et al. 1996 |
| Phi96100 | 2 | Fwd-AGGAGGACCCCAACTCCTG<br>Rev-TTGCACGAGCCATCGTAT | Chin et al. 1996 |
| phi053 | 3 | Fwd-AACCCAACGTACTCCGGCAG<br>Rev-CTGCCTCTCAGATTCAGAGATTGAC | Chin et al. 1996 |
| phi072 | 4 | Fwd-ACCGTGCATGATTAATTTCTCCAGCCTT<br>Rev-GACAGCGCGCAAATGGATTGAACT | Chin et al. 1996; Senior et al. 1996 |
| phi109188 | 5 | Fwd-AAGCTCAGAAGCCGGAGC<br>Rev-GGTCATCAAGCTCTCTGATCG | Chin et al. 1996; Senior et al. 1996 |
| phi034 | 7 | Fwd-TAGCGACAGGATGGCCTCTTCT<br>Rev-GGGGAGCACGCCTTCGTTCT | Chin et al. 1996; Senior et al. 1996 |
| phi015 | 8 | Fwd-GCAACGTACCGTACCTTTCCGA<br>Rev-ACGCTGCATTCAATTACCGGGAAG | Chin et al. 1996; Senior et al. 1996 |
| phi033 | 9 | Fwd-ATCGAAATGCAGGCGATGGTTCTC<br>Rev-ATCGAGATGTTCTACGCCCTGAAGT | Chin et al. 1996; Senior et al. 1996 |
| phi96342 | 10 | Fwd-GTAATCCCACGTCCTATCAGCC<br>Rev-TCCAACTTGAACGAACTCCTC | Chin et al. 1996; Senior et al. 1996 |
| phi427913 | 1 | Fwd-CAAAAGCTAGTCGGGGTCA<br>Rev-ATTGTTCGATGACACACTACGC | Senior et al. 1996 |

| | | | |
|---|---|---|---|
| phi127 | 2 | Fwd-ATATGCATTGCCTGGAACTGGAAGGA<br>Rev-AATTCAAACACGCCTCCCGAGTGT | Senior et al. 1996 |
| Phi051 | 7 | Fwd-CGACATCGTCAGATTATATTGCAGACCA<br>Rev-GGCGAAAGCGAACGACAACAATCTT | Senior et al. 1996 |
| phi159819 | 6 | Fwd-GATGGGCCCTAGACCAGCTT<br>Rev-GCCTCTCCCATCTCT | Senior et al. 1996 |
| phi093 | 4 | Fwd-AGTGCGTCAGCTTCATCGCCTACAAG<br>Rev-AGGCCATGCATGCTTGCAACAATGGATACA | Chin et al. 1996 |
| bnlg1194 | 8 | Fwd-GCGTTATTAAGGCAAGCTGC<br>Rev-ACGTGAAGCAGAGGATCCAT | Chin et al. 1996 |
| umc1141 | 8 | Fwd-AGAGGAGAAAGAGACAGACAGGCA<br>Rev-CAGGAACTGAATGAAAGCAACTCA | Chin et al. 1996 |
| nc004 | 4 | Fwd-TGCGAAGAAGCAGTAGCAAA<br>Rev-TGGAGGTAGAAGACGCACG | Chin et al. 1996; Senior et al. 1998 |
| bnlg1018 | 2 | Fwd-CGAGGTTAGCACCGACAAAT<br>Rev-CGAGTAAATGCTCTGTGCCA | Chin et al. 1996; Senior et al. 1998 |
| umc1930 | 10 | Fwd-TCTTCTCCAAGTGTGTTAATGCCC<br>Rev-CACACAGTGAGTCGTTTCTTTCGT | Chin et al. 1996; Senior et al. 1998 |
| phi037 | 1 | Fwd-CCCAGCTCCTGTTGTCGGCTCAGAC<br>Rev-TCCAGATCCGCCGCACCTCACGTCA | Senior et al. 1998 |

| | | | |
|---|---|---|---|
| bnlg1523 | 3 | Fwd-GAGCACAGCTAGGCAAAAGG<br><br>Rev-CTCGCACGCTCTCTCTTCTT | Senior et al. 1998 |
